# Automated Registration and Clustering for Enhanced Localization Atomic Force Microscopy of Flexible Membrane Proteins

**DOI:** 10.1101/2025.06.30.662259

**Authors:** Creighton M. Lisowski, Gavin M. King, Ioan Kosztin

**Affiliations:** Department of Physics & Astronomy, University of Missouri, Columbia, MO, USA; Department of Biochemistry, University of Missouri, Columbia, MO, USA

## Abstract

Atomic Force Microscopy (AFM) can create images of biomolecules under near-native conditions but suffers from limited lateral resolution due to the finite AFM tip size and recording frequency. The recently developed Localization Atomic Force Microscopy or LAFM (Heath et al., Nature 594, 385 (2021)) enhances lateral resolution by reconstructing peak positions in AFM image stacks, but it is less effective for flexible proteins with multiple conformations. Here we introduce an unsupervised deep learning algorithm that simultaneously registers and clusters images by protein conformation, thus making LAFM applicable to more flexible proteins. Using simulated AFM images from molecular dynamics simulations of the SecYEG translocon as a model membrane protein system, we demonstrate improved resolution for individual protein conformations. This work represents a step towards a more general LAFM algorithm that can handle biological macromolecules with multiple distinct conformational states such as SecYEG.

**Author summary:** Atomic Force Microscopy (AFM) enables high-resolution imaging of biomolecules under near-native conditions but faces lateral resolution limits due to the finite AFM tip size and recording frequency. The recently developed Localization Atomic Force Microscopy (LAFM) method addresses this by reconstructing peak positions from AFM image stacks, achieving almost atomic resolution for rigid proteins like bacteriorhodopsin (Heath et al., Nature 594, 385 (2021)). However, flexible membrane proteins with dynamic conformations, such as the SecYEG translocon, which exhibits large and highly mobile cytoplasmic loops, lead to non-physical smearing in standard LAFM reconstructions. Here, we present a computational framework combining unsupervised deep clustering and LAFM to enhance the lateral resolution of AFM images of flexible membrane proteins. Our neural network algorithm (i) groups AFM images into conformationally homogeneous clusters and (ii) registers images within each cluster. Applying LAFM separately to these clusters minimizes smearing artifacts, yielding high-resolution reconstructions for distinct conformations. We validate this approach using synthetic AFM images generated from all-atom molecular dynamics simulations of SecYEG in a solvated POPE lipid bilayer. This advancement extends LAFM’s utility to encompass conformationally diverse membrane proteins.

## Introduction

Atomic Force Microscopy (AFM) is a widely used imaging technique for studying membrane protein systems at the single-molecule level in near-native conditions. AFM images are topographic maps generated by scanning surfaces with a sharp probe, achieving near-atomic resolution in the vertical (Z) direction on many biomolecular systems [1, 2]. However, the lateral resolution (in the X-Y plane) of these images is constrained by the AFM tip geometry and size as well as the sampling discretization (recording frequency). To address these limitations, Localization Atomic Force Microscopy (LAFM) [3] has emerged as a promising post-acquisition image reconstruction method that enhances in-plane resolution.

LAFM enhances the lateral resolution of conventional AFM images by computationally reconstructing molecular details from a stack of multiple AFM images. In standard AFM imaging, lateral resolution is limited by the finite size of the scanning tip, causing structural details smaller than several nanometers to appear blurred. LAFM overcomes this limitation by exploiting subtle positional fluctuations of surface features across multiple registered AFM images. Specifically, it identifies and precisely localizes peaks corresponding to protein substructures in each individual image within the stack. By statistically combining these localized positions, LAFM reconstructs a higher-resolution image that reveals structural details previously obscured by tip geometry and sampling constraints. This approach significantly improves lateral resolution, enabling visualization of molecular conformations and sub-nanometer features that are inaccessible with traditional AFM analysis.

While effective for rigid systems (with a single stable conformation) like bacteriorhodopsin and aquaporin channel proteins [3], the direct application of LAFM to flexible proteins that undergo significant conformational changes is problematic. Indeed, in spite of its enhanced lateral resolution, the LAFM image obtained from a stack of AFM images corresponding to different protein conformations can be misleading as it blends distinct conformational states together. To mitigate this issue and generalize LAFM to more flexible membrane proteins, here we employ a two-pronged approach consisting of: (i) clustering AFM images based on conformational states, by employing deep learning (DL) algorithms, and (ii) applying LAFM selectively within clusters. Similar techniques have demonstrated success in the conformational clustering and classification of biomolecular complexes. For instance, CryoDRGN [4] employs an unsupervised DL algorithm to assist in reconstructing 3D density maps from cryo-EM data, effectively capturing and partitioning a continuous conformational landscape. Likewise, HORNET [5] leverages DL to resolve structures of flexible RNA molecules from AFM images. These approaches highlight the versatility and effectiveness of unsupervised DL algorithms, proving them a powerful tool within nanoscale imaging workflows.

Synthetic AFM images generated through molecular dynamics (MD) simulations offer a powerful approach to study flexible proteins under controlled conditions and enable direct registration and benchmarking [6]. Furthermore, the rapid innovation of GPU-accelerated computing allows for the calculation of synthetic AFM images at remarkable speeds [7]. The present study employs all-atom MD simulations of translocon SecYEG embedded in a POPE lipid bilayer. We show how LAFM can be extended to resolve structural details of this flexible membrane protein by combining LAFM methodology with clustering techniques.

## Methods

### Molecular Modeling and MD Simulation

To generate a large stack of synthetic AFM images, we performed ∼ 1*µ*s all-atom MD simulation of the protein-conduction channel SecYEG, embedded in a fully solvated 1-palmitoyl-2-oleoyl-phosphatidylethanolamine (POPE) lipid bilayer. The universally conserved SecYEG translocon is a heterotrimeric complex (composed of subunits SecY, SecE, and SecG) that plays a crucial role in mediating the transport of newly synthesized proteins either into or across the bacterial cytoplasmic membrane. The translocon SecYEG was chosen as a model membrane protein for this study because it exhibits large membrane-external loops on the cytoplasmic side of the membrane that are highly dynamic and give rise to distinct conformational states [2, 8, 9].

The crystal structure of SecYEG (PDB code 3DIN [10]), along with its orientation in the lipid bilayer, was obtained from the OPM database [11]. This structure originally contained both SecYEG and SecA, but we removed all SecA atoms to focus solely on the SecYEG complex. The CHARMM-GUI membrane builder [12–14] was employed to assemble a protein-lipid system, adding acetyl (ACE) and amide (CT2) caps to the protein termini. A homogeneous POPE lipid bilayer, representing about 75% of the native *E. coli* membrane [15], was selected to embed SecYEG while avoiding the complexity of modeling a heterogeneous bilayer. The system was subsequently solvated in a 0.03 M NaCl aqueous solution with 3 nm water padding above and below the membrane.

Next, the SecYEG-POPE system was subjected to all-atom MD simulation using NAMD2 [16], employing the CHARMM36(m) force field [17, 18]. Energy minimization and equilibration were carried out in multiple stages, gradually relaxing the restraints on the system in accordance with standard protocols [19]. A production run of unbiased MD simulation was performed at constant pressure (1 atm) and temperature (300 K), regulated by a Langevin piston barostat [20] and a Langevin thermostat [21], respectively. Atomic coordinates were saved every 0.1 ns over 1.2 *µ*s simulation, generating 12, 000 frames. To ensure the system was well equilibrated, the first 2, 000 frames were discarded, leaving 1 *µ*s of data (10, 000 frames) for analysis.

During the simulation, SecYEG displayed translational and rotational drift along the membrane surface. To facilitate analysis, we created two trajectory versions: an unaligned (original) trajectory and an aligned trajectory. For the aligned version, using VMD [22], we minimized the root mean squared displacement (RMSD) of transmembrane helix backbone atoms relative to their initial positions. This alignment was essential since subsequent analyses rely on consistent protein positioning, with the aligned trajectory serving as our reference standard for comparisons.

### Simulated AFM Images

Simulated AFM (SimAFM) images were generated from all-atom MD trajectories (see Fig. 1) using our open-source Python package, AFMpy [23]. The SimAFM implementation in AFMpy employs the MDAnalysis [24, 25] library for trajectory processing, NumPy [26] for efficient vectorized calculation of tip contacts, and CuPy [27] for GPU acceleration.

**Fig 1.**
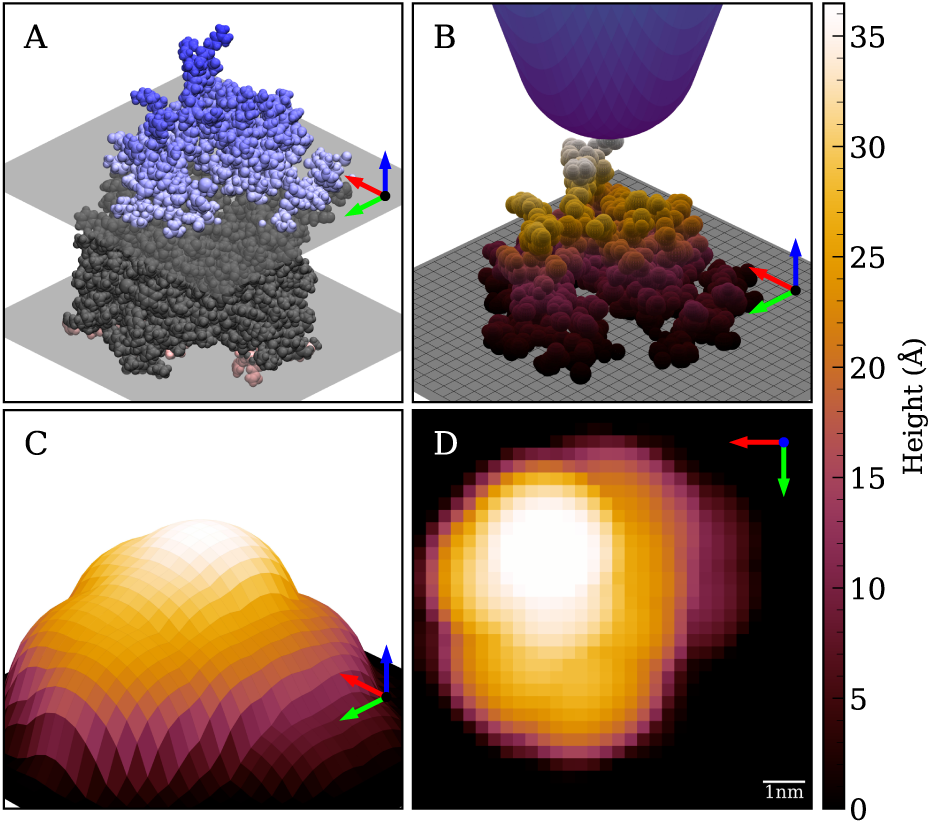
Simulated Atomic Force Microscopy (SimAFM) of SecYEG. **A.** Van der Waals representation of SecYEG rendered in VMD [22]. Average lipid head group positions and transmembrane protein regions are shown in gray, while only the cytoplasmic (blue) and periplasmic (red) protrusions of the protein are accessible to the AFM tip during scanning. **B.** Simulated AFM tip (blue) contacting the cytoplasmic side of SecYEG, with tip contact calculated at each grid point across the *X*-*Y* plane. **C.** Surface representation of the SimAFM scan, showing how molecular details are blurred by the finite geometry of the AFM tip. **D.** Top-down view of the SimAFM scan of the cytoplasmic side of SecYEG.

Each frame from the MD trajectory is loaded and rotated so that the membrane side of interest faces the +*Z* direction. The membrane plane is defined by the average *Z*-coordinate of the phosphorus atoms in the upper leaflet, which serves as the background. Protein atoms located above this background are represented as solid spheres with their van der Waals radii, as specified by the CHARMM force field [28]. The *X*–*Y* plane is then divided into a uniform two-dimensional grid according to the chosen scan boundaries and pixel resolution.

The simulated AFM tip is modeled as a cone with a spherical apex, characterized by its radius (*R*) and a half-angle *θ* = 18*^◦^*, matching the typical value for experimental AFM tips. To generate the SimAFM image, the tip is virtually scanned across each point of the grid, and the height at which it contacts the protein surface is recorded. This procedure is repeated for every frame in the trajectory, resulting in a stack of SimAFM images.

Stacks of SimAFM images were generated to visualize the protruding regions of SecYEG on both the cytoplasmic and periplasmic sides of the membrane, using both aligned and unaligned MD trajectories of the SecYEG-POPE system. To mimic a range of experimentally available AFM tips, the simulated tip radius was varied, including an idealized tip with *R* = 2Å. Scan resolution was also systematically adjusted, producing SimAFM stacks across a broad spectrum of resolutions. Specifically, tip radii from 2Å to 80Å and scan resolutions from 2Å/px to 10Å/px were used.

Stacks were assigned a code reflecting their alignment status (Aligned: A; Unaligned: U), membrane side (Cytoplasmic: C; Periplasmic: P), tip radius (in Å), and scan resolution (in Å/px). For instance, a stack of aligned cytoplasmic images simulated with a 20Å tip and a 2Å/px scan resolution is labeled as AC–20–2 for easy identification. Notable stacks include those generated with tip radii of 2Å (approximating the width of a carbon atom), 20Å (representing the PeakForce-HIRS-F-B tip), and 80Å (representing the BioLever Mini AC40 tip).

### Localization Atomic Force Microscopy (LAFM)

Each stack of images was processed to generate an enhanced lateral resolution image using our Python implementation of the LAFM algorithm [3]. The workflow consists of several steps: First, using OpenCV [29], the input image stack is enhanced to a higher, user-defined resolution by applying bicubic interpolation to expand each image. Next, peak detection is performed on each interpolated image to identify local maxima, which are then broadened using a Gaussian kernel of user-specified width, producing images where each peak is represented by a 2D Gaussian. The height of each Gaussian is scaled between 0 and 1 according to the detected real-space height relative to the maximum in the stack. These Gaussian images are then averaged pixel-wise across the entire stack to create a cumulative peaking probability image. Finally, this probability image is multiplied by the per-pixel averaged real-space height image to produce the final LAFM image.

### Structural Similarity Index Measure (SSIM)

The quality of LAFM images relative to the benchmark (i.e., the LAFM images AP–2–2 and AC–2–2, respectively) was evaluated using the Structural Similarity Index Measure (SSIM) [30], which offers several advantages over simpler metrics such as mean squared error [31]. SSIM is particularly well-suited for this application because its focus on local structural similarity more closely reflects perceived visual quality. However, when applying SSIM to protein AFM images, special care is required: the uniform 0-height background in LAFM images creates large areas of perfect similarity, which can bias the SSIM calculation. To address this, a masked SSIM approach was used. A mask was generated by selecting pixels above a threshold set at 5% of the image’s maximum height, effectively isolating the protein features from the background. SSIM was then computed over the entire image, and the mask was applied to exclude background pixels. The mean value of the masked SSIM image was reported as the final similarity score.

LAFM images generated from each aligned simulation stack were compared to the benchmark using the masked SSIM score, with the results recorded in 2D arrays called LAFM quality profiles. These profiles are visualized as topographic maps (see Fig. 5), illustrating LAFM quality as a function of tip radius (*R*) and scanning resolution (*P*).

### Deep Spectral Clustering (DSC)

Given an aligned stack of images of a flexible protein exhibiting distinct structural conformations, it is necessary to first identify and separate (i.e., cluster) these conformations before applying the LAFM algorithm. In AFMpy, this conformational clustering is performed using a deep spectral clustering (DSC) algorithm, as illustrated in Fig. 2. Our DSC implementation uses the Spectral Clustering module from scikit-learn [32] and employs the Keras API in TensorFlow [33, 34] for constructing, training, and validating the deep learning models.

**Fig 2.**
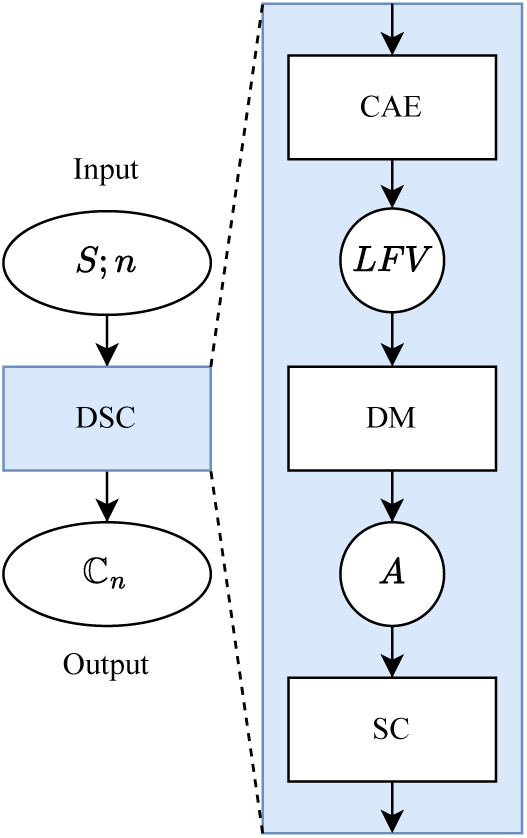
Deep Spectral Clustering (DSC) Algorithm: A convolutional autoencoder (CAE) is trained on input image stack (*S*) to extract latent feature vectors (*LFV*). *LFV* pairs are compared by a given distance metric (DM) to construct an affinity matrix (*A*). Spectral clustering (SC) is applied to *A*, returning *n* cluster labels for the input stack. Images are grouped by label into clusters (*C*) which make up the set of *n* clusters (ℂ_*n*_ = {*C*_1_*, C*_2_*, . . . , C*_*n*_}).

DSC assigns *n* conformational cluster labels to a stack of images by applying spectral clustering to latent feature vectors (LFVs) extracted by a convolutional autoencoder (CAE) [35]. The CAE is first trained end-to-end to minimize a reconstruction loss, defined as 1 − similarity metric, where SSIM is used as the similarity metric due to its effectiveness in capturing protein structural features. To properly account for both protein and background pixels, a weighted SSIM loss is employed, combining standard SSIM (to address background) and masked SSIM (to focus on protein features). After training, LFVs are extracted from the images, and an affinity matrix (*A*) is constructed by pairwise comparison of these LFVs using locally scaled affinity [36]. Spectral clustering is then performed on this matrix to assign cluster labels to the images.

Because the SimAFM stack captures a continuous range of motion, it contains both images of stable conformations and transient states during transitions. To separate these conformations, DSC was applied in a hierarchical manner. First, *n* = 2 DSC is applied to the stack, and the resulting clustered LAFM images are generated. If the clustered LAFM images have high similarity, i.e., the masked SSIM exceeds a set threshold, the clusters are combined, and the process halts. If the two LAFM images differ, however, the affinity matrix is reclustered for *n* = 3, and new clustered LAFM images are generated. These *n* = 3 clustered LAFM images are cross-compared with the *n* = 2 LAFM images. Images with high similarity represent stable conformations, and the *n* = 3 labeled images are removed from the stack. This process iterates, using the reduced stack and affinity matrix as input, until no stable clusters are found, or the number of remaining unclustered images falls below a set minimum cluster size. We refer to this process as Hierarchical DSC (HDSC).

### Registration and Clustering (REC)

Given an unaligned image stack, conformational clustering can be performed using the registration and clustering (REC) algorithm as outlined in Fig. 3. The REC algorithm operates in two modes depending on the number of provided registration references. With a single reference and a specified number of clusters *n*, the algorithm registers all images to the reference and applies DSC to identify *n* registered clusters. Image registration was performed using the pystackreg Python package [37], applying rigid body transformations (translation and rotation only). This ensured alignment of images without scaling or shearing. If *n* registration references are provided, the stack is registered separately to each reference, generating *n* registered stacks. Each stack is then clustered independently using DSC, resulting in a superset of registered clusters. Clusters containing their respective registration reference images are considered well-registered and are retained for the final output. Additionally, within each cluster, the image with the highest silhouette score [38] is selected as an improved registration reference, yielding a refined set of registration references.

**Fig 3.**
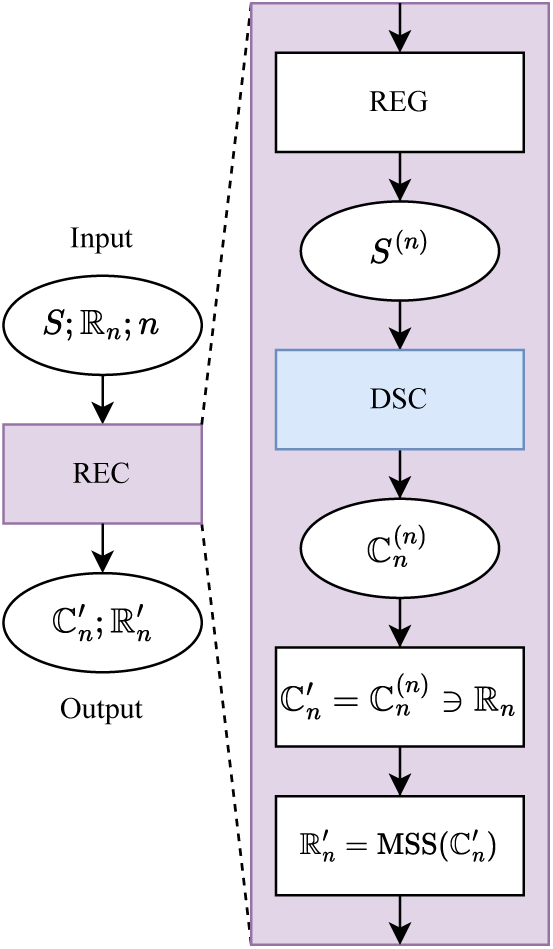
Registration and Clustering (REC) Algorithm: The input image stack (*S*) is registered (REG) using each image in the reference set ℝ_*n*_ = {*R*_1_*, R*_2_*, . . . , R*_*n*_}, resulting in *n* registered stacks (*S*^(*n*)^). Each registered stack is then subjected to DSC clustering, producing a set of clusters (ℂ_*n*_) for each reference. These cluster sets are combined into a registered cluster superset 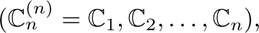 which contains the clusters generated for each registration reference. From this superset, only the clusters corresponding to their respective registration references 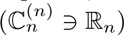 are retained to form the refined cluster set 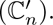 Additionally, for each cluster, the image with the maximum *silhouette score* (MSS) is chosen to create a set of refined registration references 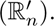

### Iterative Registration and Clustering (IREC)

Selecting appropriate registration references and determining the optimal number of clusters are challenging tasks. To address this, we developed an iterative registration and clustering (IREC) algorithm. As outlined in Fig. 4, IREC iteratively refines both cluster assignments and registration references to simultaneously optimize the registration references and the number of clusters present in a conformationally diverse AFM image stack. The IREC implementation leverages registration methods from the Python module pyStackReg [39] and clustering metrics from scikit-learn [32].

**Fig 4.**
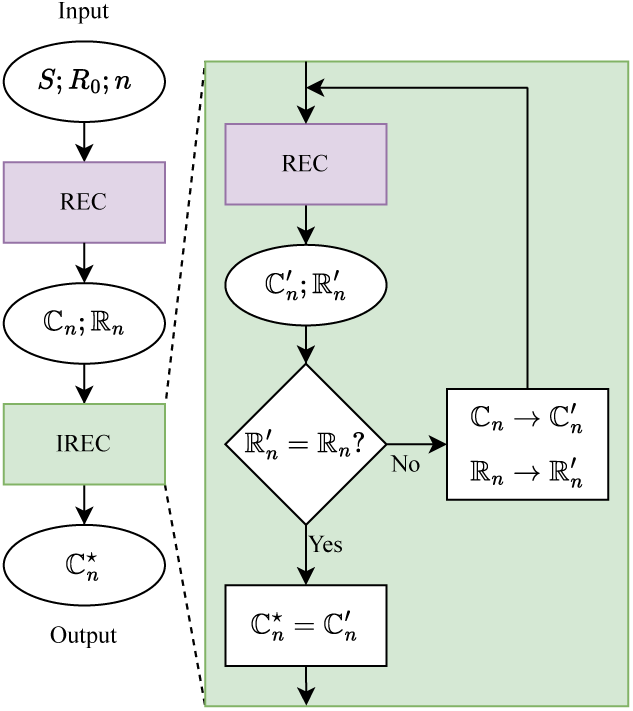
Iterative Registration and Clustering (IREC) Algorithm: The input image stack (*S*) is first aligned and grouped using registration and clustering (REC) based on an initial reference image (*R*_0_). This yields an initial set of clusters (ℂ_*n*_) and corresponding registration references (ℝ_*n*_), which are then iteratively refined. In each iteration, every reference image in ℝ_*n*_ is used to re-register and re-cluster *S* via REC, producing updated clusters 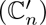 and new references 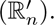 The new references are compared to those from the previous iteration. If they differ 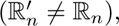 the references are updated 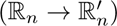 and the process repeats. Iteration continues until the references converge 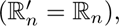 a cyclic pattern is detected, or a maximum number of iterations is reached, at which point the algorithm outputs the final, optimized cluster set 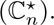

For a given *n*, the input image stack (*S*) is first processed by REC using an initial reference (*R*_0_), producing *n* clusters (ℂ_*n*_) and their corresponding references (ℝ_*n*_). These clusters and references are then refined through iterative cycles: in each iteration, the stack is re-registered and re-clustered according to the refined references from the previous step, yielding new clusters and references. The newly generated references are compared to those from the preceding iteration; if discrepancies are found, the mismatched references are updated and the process repeats. Iteration continues until all references match, cyclic behavior is detected, or a maximum number of iterations is reached, at which point the clusters are considered optimal 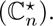 Finally, LAFM is applied to each optimal cluster to generate the set of enhanced images.

Although IREC determines the optimal conformational clusters for a given *n*, it remains uncertain whether *n* is the optimal number of clusters required to partition the image stack. To assess the appropriate cluster count, the clustered LAFM images are compared as *n* increases. First, IREC is applied to the image stack to generate the corresponding LAFM images. Since these LAFM images are generated using different registration references, they may not be mutually aligned. To evaluate their similarity, the LAFM images are registered to one another and compared using masked SSIM. If all SSIM scores fall below a predefined similarity threshold (e.g., 0.8), the enhanced images are considered to represent distinct conformations, indicating that at least *n* clusters are necessary. However, this test does not exclude the possibility that more clusters may be required, so IREC is reapplied with *n* incremented by one. This process is repeated for increasing values of *n* until any SSIM score between clustered LAFM images exceeds the similarity threshold, indicating that the optimal number of clusters for the image stack is *n* − 1.

### Computational Performance

Benchmark computations were performed to assess the acceleration and scalability of AFMpy functions across CPU and GPU processing. All benchmarks ran on a Linux workstation with an Intel Xeon CPU (w9–3495X, with 56 cores) and two NVIDIA RTX A5500 GPUs (connected via NVLink). Table 1 summarizes execution times for SimAFM, DSC, Hierarchical DSC, and IREC in both CPU-only and GPU-enabled modes.

**Table 1.** CPU vs. GPU runtime and speedup.

| Process | CPU Time (s) | GPU Time (s) | GPU Speedup |
| --- | --- | --- | --- |
| Periplasmic SimAFM | 43 | 31 | 1.4 |
| Cytoplasmic SimAFM | 344 | 68 | 5.1 |
| DSC | 376 | 33 | 11.5 |
| Hierarchical DSC | 392 | 50 | 7.9 |
| IREC | 4598 | 493 | 9.3 |

The 10,000-frame stacks of 32×32 simulated AFM images for periplasmic (110 protruding atoms) and cytoplasmic SecYEG (∼2000 protruding atoms) were generated using SimAFM from the 1 *µ*s MD trajectory. Simulation time increased with system size; GPU acceleration provided greater speedup for the larger cytoplasmic SecYEG.

Benchmarking of DSC, Hierarchical DSC, and IREC functions on the cytoplasmic SecYEG stack showed that GPU acceleration substantially reduced computation times for all deep learning-based analyses, with speedups of 7.9 − 11.5. These functions employ TensorFlow (GPU-accelerated) and scikit-learn (CPU-based) components. Based on the obtained results, we recommend prioritizing GPU resources for AFMpy workflows, especially for deep learning components and SimAFM simulations of large systems or long trajectories.

## Results and Discussion

### LAFM Quality Assessment

To quantify the influence of the AFM tip radius (*R*) and pixel resolution (*P*) on the quality of LAFM images, we compared images derived from the aligned cytoplasmic and periplasmic SecYEG SimAFM stacks to their respective benchmark LAFMs (from the AC–2–2 and AP–2–2 stacks) using masked SSIM. The results are shown as density plots in the left panel of Fig. 5. These LAFM quality profiles, for both periplasmic and cytoplasmic SecYEG, show that the LAFM image quality increases nonlinearly with decreasing *R* and *P*, though with notable differences between the two faces. For periplasmic LAFM, achieving a “good” quality threshold of 0.7 SSIM requires at least an *R* = 4 nm tip scanning at *P* = 4 Å/px, indicating that both parameters are important, and moderate tip sizes and resolutions are sufficient for high-quality reconstruction. In contrast, for the cytoplasmic side, scanning resolution has less influence on the LAFM image quality, while tip radius plays a much more significant role. Here, obtaining a “good”, 0.7 SSIM quality LAFM image requires a much finer *R* = 1 nm tip, even when scanning at the same *P* = 4 Å/px resolution. Therefore, compared to the periplasmic side, cytoplasmic LAFM images require sharper AFM tips to effectively resolve structural details at equivalent pixel resolution.

**Fig 5.**
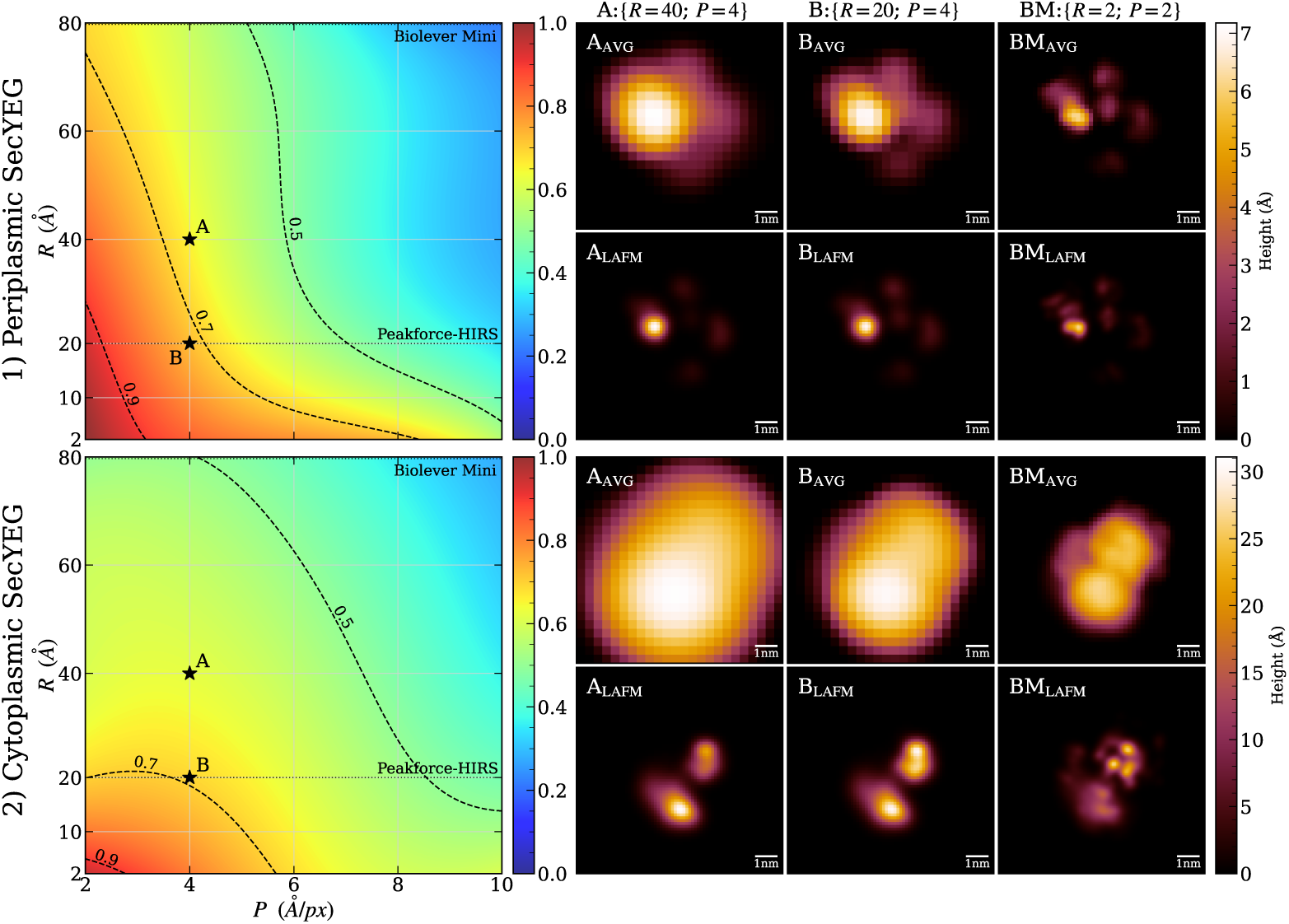
**Left:** Quality profile of LAFM images for 1) periplasmic (top) and 2) cytoplasmic (bottom) SecYEG. Color-coded density plots display masked SSIM scores of simulated LAFM images as functions of AFM tip radius (*R*) and pixel resolution (*P*), benchmarked against reference LAFM images acquired with *R* = 2 and *P* = 2. The dotted horizontal lines indicate the nominal radii of a commercial AFM tip (PeakForce-HIRS tip, *R* = 20). The radius of another common tip (Biolever Mini, *R* = 80), coincides with the top axes of the plots. Dashed contour lines at SSIM values of 0.9, 0.7, and 0.5 delineate the boundaries between “very good”, “good”, and “acceptable” LAFM quality regions. **Right:** Per-pixel averaged (AVG) AFM and LAFM images of 1) periplasmic (top) and 2) cytoplasmic (bottom) SecYEG for A:{*R* = 40; *P* = 4}, B:{*R* = 20; *P* = 4} and benchmark BM:{*R* = 2; *P* = 2} image stacks.

As shown in the right panel of Fig. 5, LAFM effectively resolves five distinct domains in periplasmic SecYEG using the AP–40–4 dataset (SSIM = 0.670) and further enhances feature accuracy in the AP–20–4 stack (SSIM = 0.748). For cytoplasmic SecYEG, the average AFM image appears as a featureless blob, whereas the corresponding LAFM image reveals three distinct domains. However, this improvement is modest: the AC–40–4 image yields a relatively low SSIM of 0.632, and reducing the tip radius to 2 nm (AC–20–4) results in only a marginal increase to 0.662.

A notable challenge in the cytoplasmic case arises from conformational changes in the approximately 35-amino acid loops connecting transmembrane helices 6-7 and 8-9 within the SecYEG structure. These dynamic regions cause the bottom domain to appear smeared upward and to the left in the LAFM image, which inherently blends multiple structural states into a single composite representation. To address this limitation, deep spectral clustering approaches will be essential for distinguishing and separating the distinct conformational states present within the image stack, thereby enabling more precise structural characterization of dynamic membrane protein complexes.

### Hierarchical Deep Spectral Clustering Results

Hierarchical DSC was applied to the cytoplasmic side images (AC–20–4), identifying four conformational clusters. The corresponding LAFM images are shown in Fig. 6. Compared to the unclustered LAFM image, where different conformations are blended together, clustering clearly separates these states, with each cluster producing a unique LAFM image representing a specific conformation. The clusters are shown in order of decreasing population. The stack is dominated by clusters A and B, which together account for 86.5% of the total, indicating that these conformations represent metastable states. In contrast, clusters C and D are less populated (collectively 13.5%), and most likely represent short-lived transition states. Furthermore, since we know the true sequence of frames, the temporal progression of cluster occupancy indicates that AC–20–4 starts in conformation A and transitions to conformation B, briefly passing through conformations C and D. However, it is important to emphasize that our DSC algorithm produces consistent results regardless of image sequence, yielding the same outcome when applied to the AFM image stack even after randomizing the order of frames.

**Fig 6.**
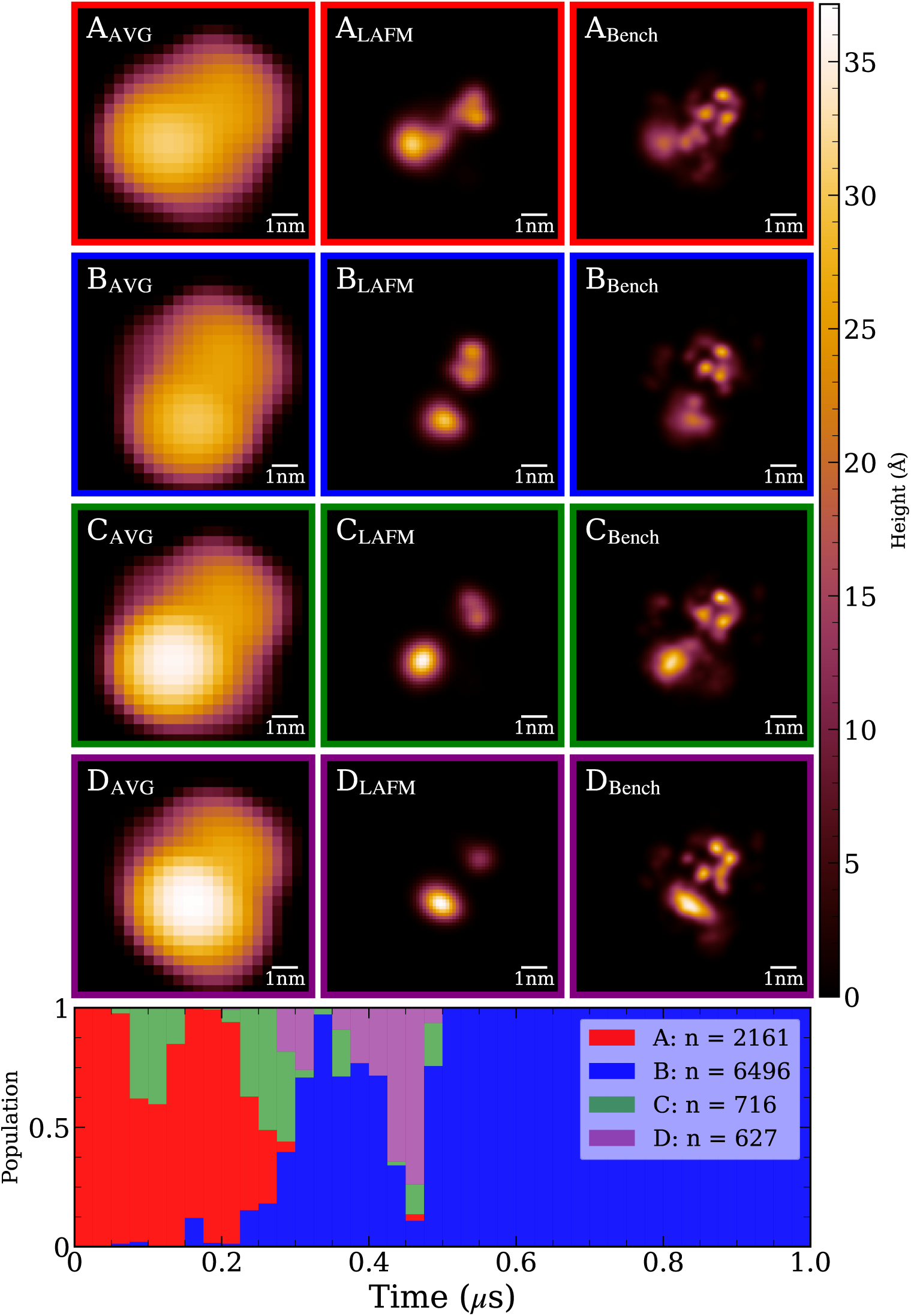
**Top:** Hierarchical deep spectral clustering results for AC–20–4. Each row shows the average, LAFM, and benchmark LAFM images for the detected conformational clusters. **Bottom:** Cluster occupancy over the trajectory, shown in discrete *t* = 0.025*µ*s bins. Despite the input frames being shuffled during clustering, the reconstructed time series displays a clear progression between states. The cytoplasmic side of SecYEG starts in conformation A. Clusters C and D, whose populations fall below the minimum stable cluster size, are considered transient states that the protein passes through before settling into the stable conformation B.

For completeness, we also applied hierarchical DSC to periplasmic side images (AP–40–4) and identified two distinct conformational clusters, as shown in Fig. 7. The resulting LAFM images display similar features in the top and leftmost regions, but the right domain is noticeably taller in conformation B, suggesting that the protruding loop extends outward, perpendicular to the membrane surface. Overall, the periplasmic side of SecYEG exhibits minimal large-scale conformational changes. Cluster A is predominant, representing 97.9% of the stack, while cluster B appears only briefly, accounting for 2.1%. Although the transition between these states is subtle, DSC reliably distinguishes them, demonstrating the algorithm’s sensitivity to minor conformational differences.

**Fig 7.**
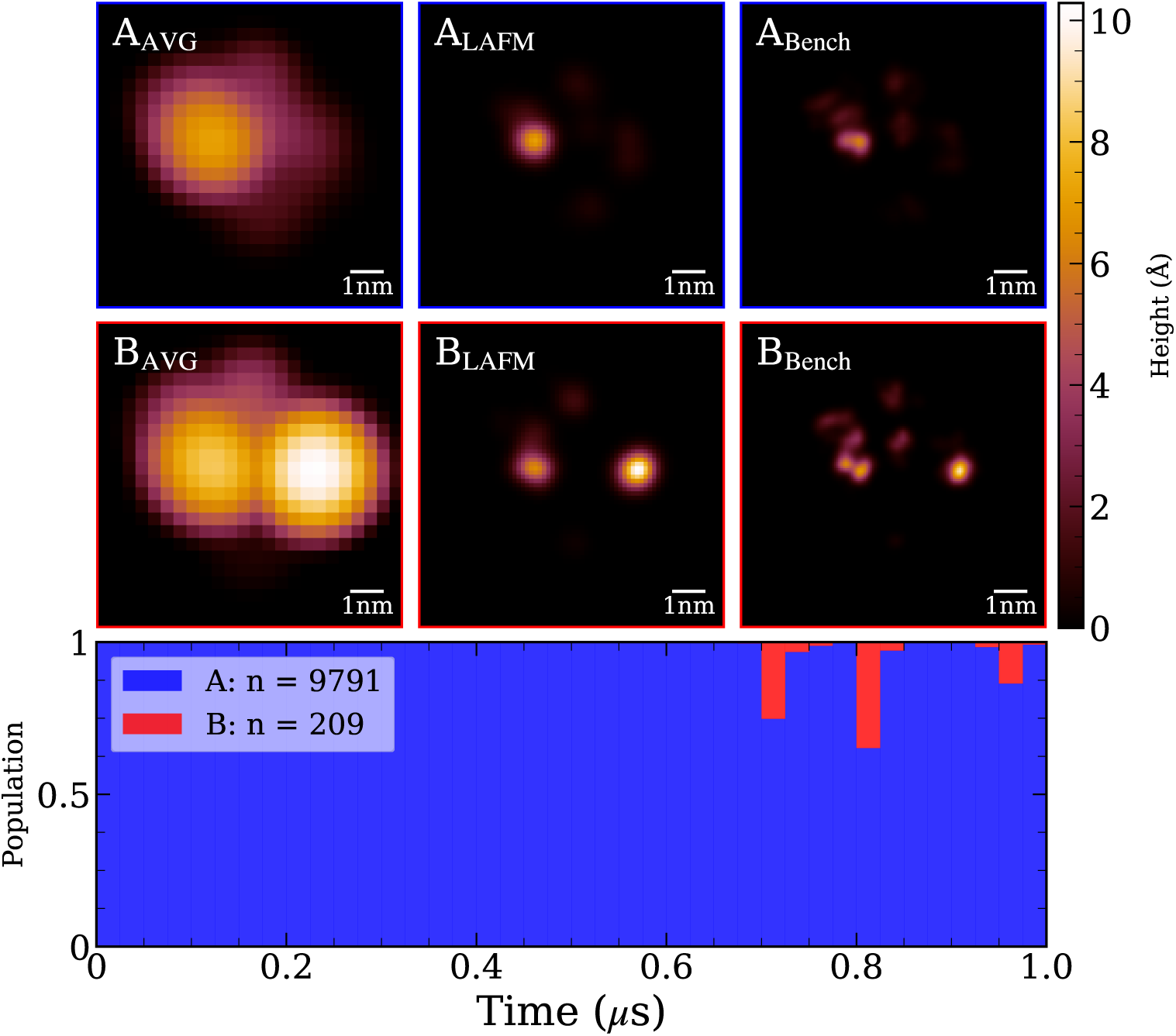
**Top:** Hierarchical deep spectral clustering results for AP–20–4. Each row shows the average, LAFM, and benchmark LAFM images for the detected conformational clusters. **Bottom:** Cluster distribution over the trajectory (binned in 0.025*µ*s intervals). The periplasmic side of SecYEG predominantly adopts the stable conformation represented by cluster A throughout most of the simulation. Around 0.7*µ*s, SecYEG briefly transitions to the conformation of cluster B, with subsequent intermittent switching between clusters A and B. Since cluster B contains fewer frames than the minimum threshold for a stable cluster, it is classified as a transient conformation.

### Iterative Registration and Clustering (IREC) Results

We applied IREC to the UC–20–4 stack, generating cytoplasmic LAFM images for increasing numbers of clusters. The results are presented in Fig. 8. The unclustered LAFM image (first row) alone cannot establish whether a single stable conformation exists; therefore, we applied IREC with *n* = 2 clusters. The resulting two-cluster LAFM images (second row) were compared using SSIM, yielding scores below the 0.9 threshold, thereby confirming that these images represent distinct conformations and that at least two clusters are required. Subsequently, we applied IREC with *n* = 3 clusters. The three resulting LAFM images (third row) again exhibited SSIM scores below 0.9, validating the presence of distinct conformations and indicating that three clusters are necessary. Finally, we tested *n* = 4 clusters. Cross-comparison of the four-cluster LAFM images (fourth row) revealed that clusters *D*_2_ and *D*_3_ shared an SSIM score of 1.0, indicating they represent identical conformations. Therefore, *n* = 3 represents the optimal clustering solution for this system.

**Fig 8.**
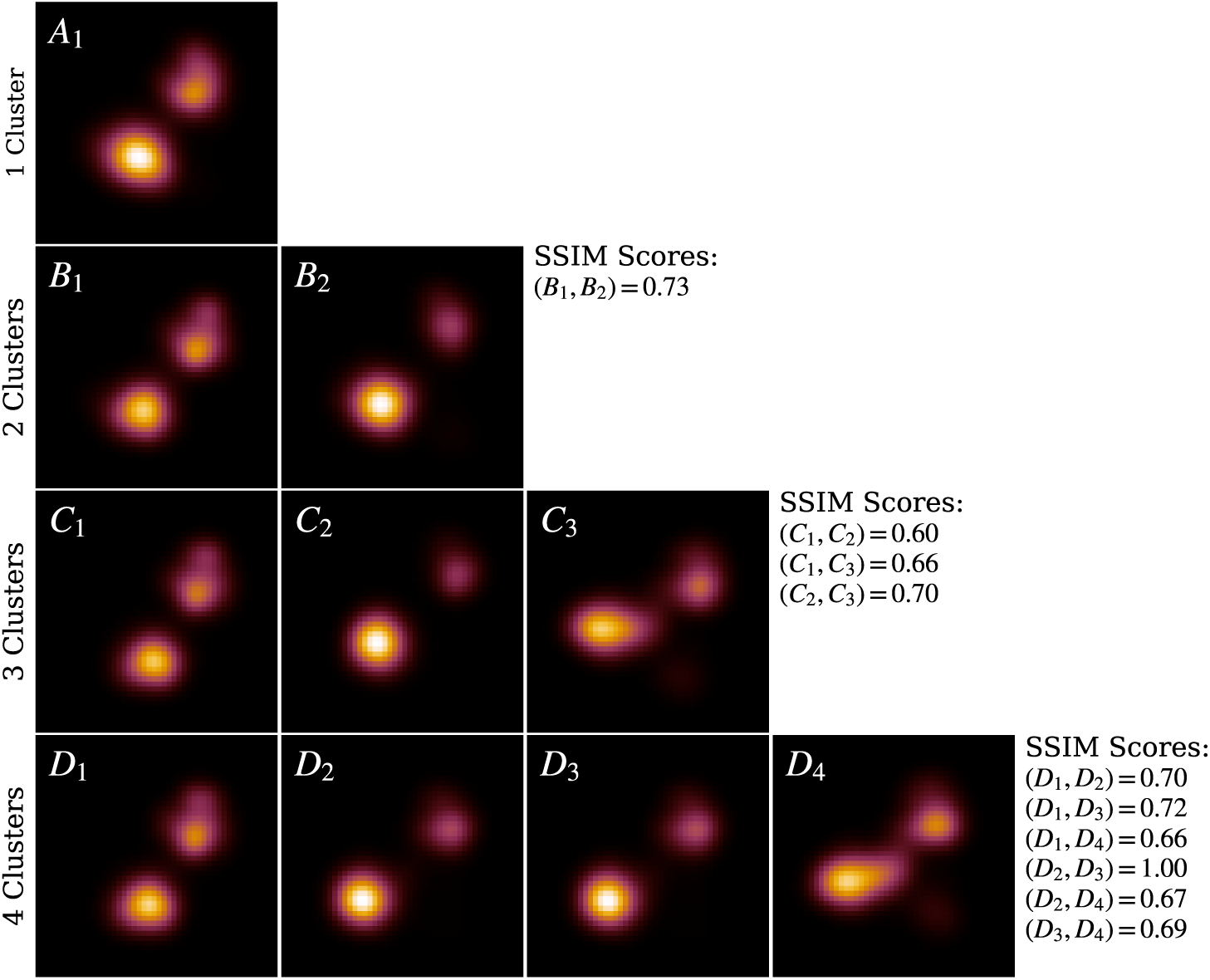
IREC applied to UC–20–4. LAFM images for each detected conformational cluster are shown, with rows corresponding to increasing numbers of clusters. For *n* = 3, the clustered LAFM images have SSIM scores below the 0.9 threshold and closely resemble those obtained from hierarchical DSC on the aligned stack. With *n* = 4, two of the clustered LAFMs exhibit a similarity score of 0.99, indicating redundancy and confirming that *n* = 3 is the optimal clustering solution.

Visual comparison of the three IREC-derived clusters with those obtained from hierarchical DSC analysis of the aligned cytoplasmic stack (Fig. 6) reveals strong correspondence: IREC cluster *C*_1_ corresponds to DSC cluster *B*, *C*_2_ to cluster *C*, and *C*_3_ to cluster *A*. Notably, all three major conformational states are successfully resolved despite the use of unaligned input data. The absence of DSC cluster *D* in the IREC analysis suggests that frames originally assigned to this cluster are redistributed among neighboring clusters during processing of the unaligned stack.

We also applied IREC to the UP–40–4 stack (Fig. 9). The first row displays the unclustered LAFM image, which lacks definitive evidence for a single, stable conformation. Consequently, we applied IREC with *n* = 2, yielding two LAFM images with an SSIM score of 0.50, indicating distinct conformational states and validating the requirement for at least two clusters. Upon increasing to *n* = 3, cross-comparison analysis revealed that clusters *C*_2_ and *C*_3_ exhibited an SSIM score of 1.0, demonstrating they represent identical conformations. Therefore, *n* = 2 constitutes the optimal clustering solution.

**Fig 9.**
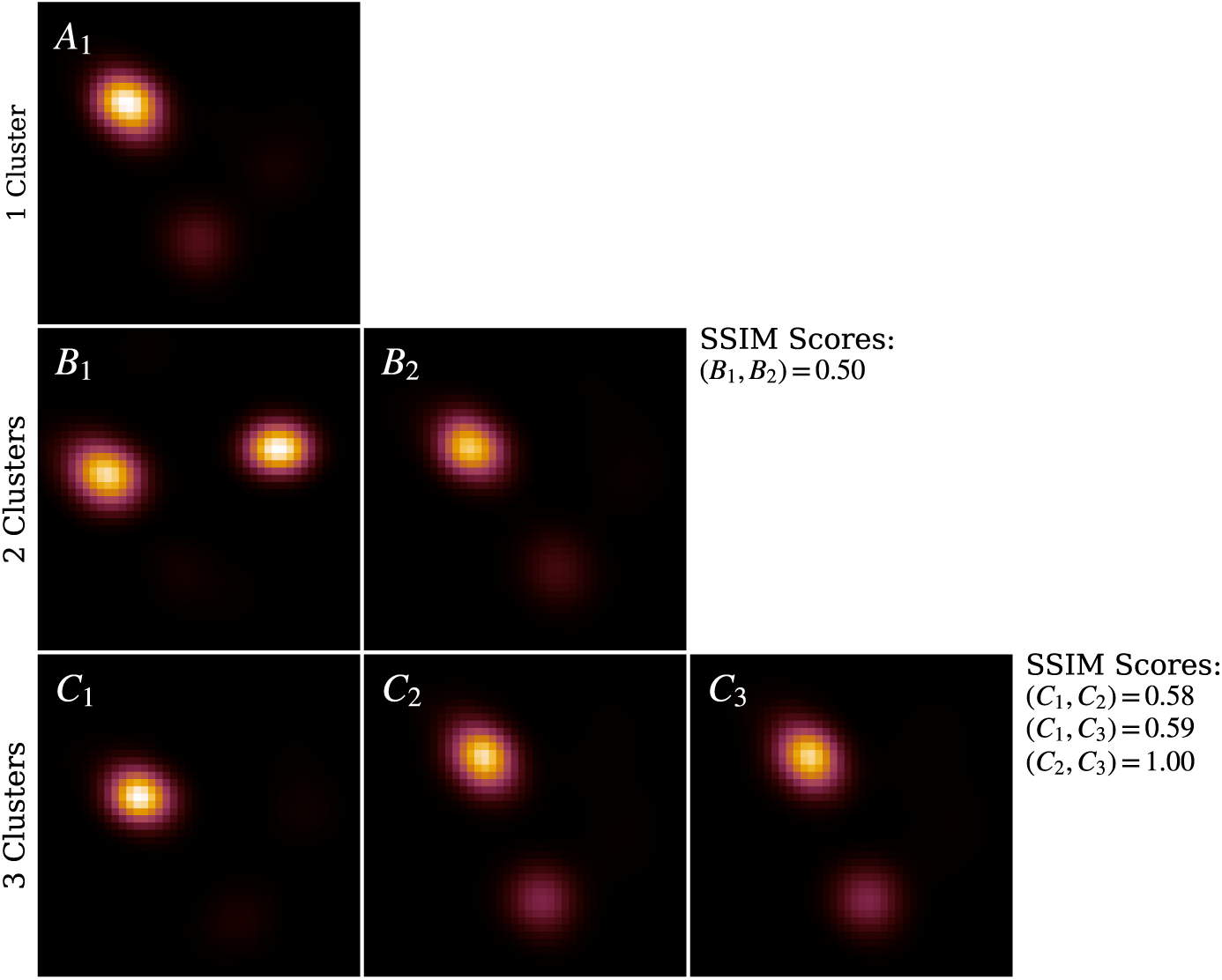
IREC applied to UP–40–4. LAFM images for each detected conformational cluster are shown, with rows corresponding to increasing numbers of clusters. For *n* = 2, the clustered LAFM images exhibit SSIM scores below the 0.9 threshold and closely resemble those obtained from hierarchical DSC on the aligned stack. With *n* = 3, two clustered LAFMs show a 0.99 SSIM score, indicating redundancy and confirming that *n* = 2 is the optimal clustering solution.

Visual comparison between these two IREC-derived clusters and the two-cluster solution obtained from hierarchical DSC (aligned stack) demonstrates that IREC cluster *B*_1_ corresponds to DSC cluster *B*, while *B*_2_ corresponds to cluster *A*. This correspondence confirms comprehensive capture of all conformational states without missing clusters, thereby validating IREC’s accuracy for the periplasmic dataset.

To further evaluate IREC, we combined the UP–20–4 and UC–20–4 stacks and applied IREC to this merged dataset (Fig. 10). With *n* = 2, IREC successfully separated cytoplasmic and periplasmic images into distinct clusters. Since the SSIM between these clusters was substantially below the 0.9 similarity threshold, we systematically increased the number of clusters to *n* = 3, which preserved the compartmental separation while subdividing the cytoplasmic images into two distinct conformational clusters. All pairwise SSIM scores remained below the established cutoff, confirming the identification of genuinely distinct conformations.

**Fig 10.**
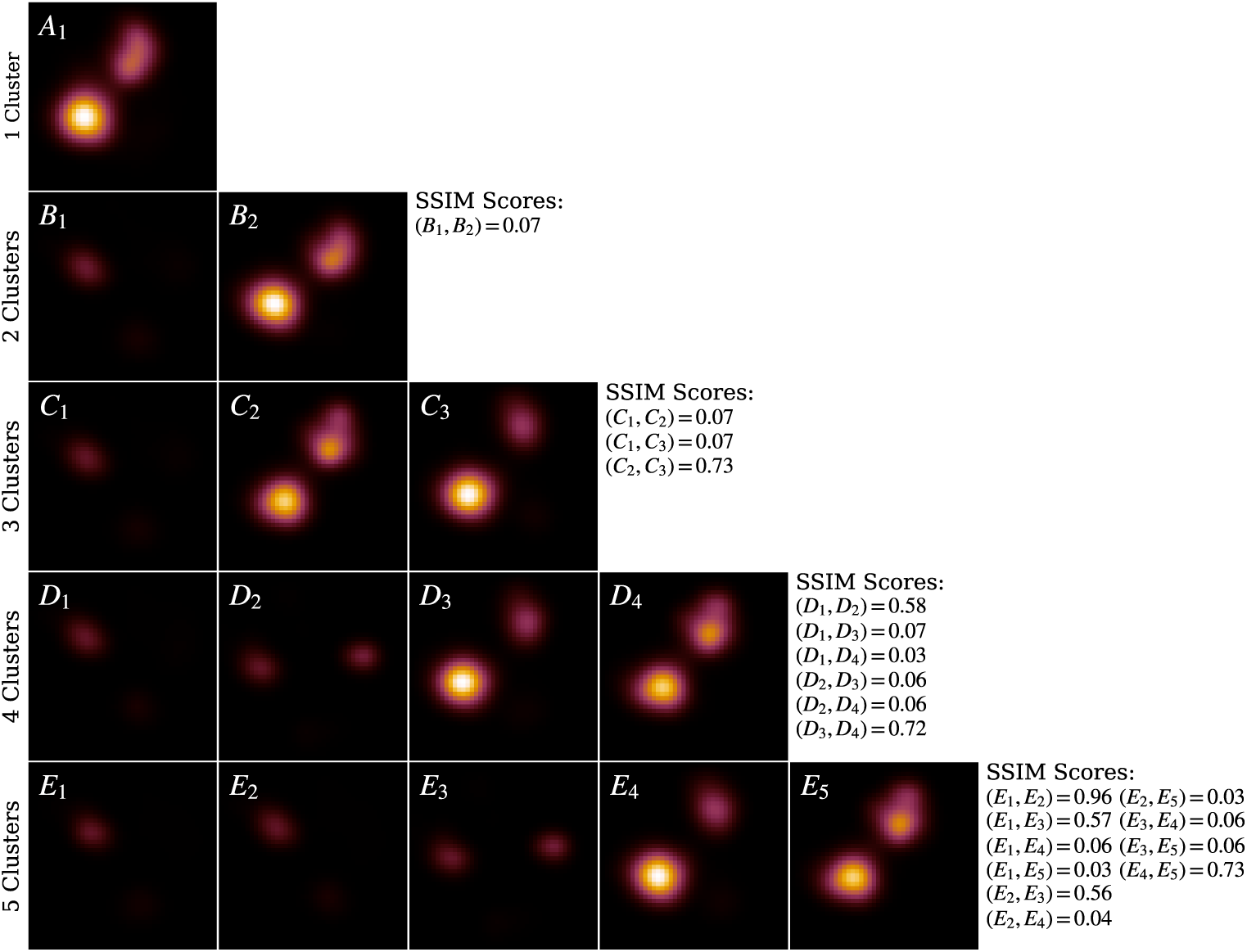
IREC applied to combined UP-20-4 and UC-20-4. LAFM images for detected conformational clusters are shown, with rows corresponding to increasing cluster numbers. At *n* = 2, clusters perfectly separate cytoplasmic and periplasmic sides, generating distinct LAFM images for each orientation. *n* = 3 retains side separation while resolving two cytoplasmic conformational clusters. *n* = 4 further splits both cytoplasmic and periplasmic orientations into two clusters each. *n* = 5 reproduces existing clusters but duplicates a periplasmic conformation, confirming *n* = 4 as the optimal clustering solution.

We extended this analysis to *n* = 4 and *n* = 5 clusters. At *n* = 5, two clusters exhibited an SSIM of 0.96, exceeding our similarity threshold and indicating they likely represent the same conformational state. Therefore, *n* = 4 was determined to be the optimal clustering solution, providing the maximum resolution of distinct conformational states without over-partitioning the data.

This approach successfully identified two periplasmic and two cytoplasmic clusters. However, the third cytoplasmic conformation observed in the UC–20–4 dataset was not detected. Analysis of the initial references employed by REC revealed an inherent bias: the algorithm utilized two cytoplasmic and three periplasmic references, which during iterative refinement failed to converge on the missing cytoplasmic conformation. These findings indicate that when analyzing datasets containing structurally distinct morphologies (e.g., periplasmic versus cytoplasmic conformations), IREC should be applied hierarchically — first partitioning the data by major structural class, then performing separate clustering within each subset to resolve finer conformational distinctions.

## Conclusion

The full power of the atomic force microscope – directly visualizing individual atoms – is typically realized only when imaging extremely flat and inert solid-state surfaces such as silicon or mica. The development of LAFM, which enables the distinction of individual amino acid residues in hydrated protein specimens, marks a significant advance for force microscopy in biophysical research. However, the application of LAFM has thus far been largely restricted to conformationally stable proteins with relatively rigid structures.

This work represents a significant step toward extending LAFM to a broader class of biological macromolecules that exhibit greater conformational flexibility and populate multiple structural states. Leveraging the AFMpy Python package [23] developed here, we combined LAFM with deep learning algorithms to both identify and enhance the lateral resolution of individual protein conformations. Using the core component of the general secretory system of *E. coli*, the translocon SecYEG, as a model system, we demonstrated the feasibility of this approach on simulated AFM image stacks.

While our results are based on simulated AFM images over computationally accessible timescales, this work lays the foundation for future studies employing experimental data, such as high-speed AFM (HS-AFM) images, to probe protein conformational landscapes over biologically relevant timescales. Ultimately, these advances may enable the routine application of LAFM to a wider array of dynamic biomolecular systems.

## Acknowledgments

This work was supported in part by the National Science Foundation (NSF) under grant number 2122027.

